# Molecular model of the nuclear pore complex reveals a thermoreversible FG-network with distinct territories occupied by different FG motifs

**DOI:** 10.1101/568865

**Authors:** Kai Huang, Mario Tagliazucchi, Sung Hyun Park, Yitzhak Rabin, Igal Szleifer

## Abstract

Despite the intensive study of the nuclear pore complex (NPC), its functional core, the central transporter, remains poorly understood. Here, we investigate this unfolded and dynamic part of the NPC using a molecular theory that addresses both entropic and enthalpic effects of the intrinsically disordered phenylalanine-glycine-rich nucleoporins (FG-Nups). Our model shows that the cooperative effect of FG-pairing, specific spacer cohesion, and charge interaction leads to a remarkably elaborate gating structure inside the NPC. In particular, we find sequence-programmed “phase separation” between charge-rich and charge-poor regions, and a polarized electrostatic potential throughout the NPC. The model predicts a thermoreversible FG-network with inhomogeneous FG-pairing fraction in space, which features distinct territories of different types of FG motifs. Our theoretical anatomy of the central transporter reveals a clear sequence-structure-function relationship of the FG-Nups, and explains in a self-consistent way how nucleocytoplasmic transport can be efficient yet specific.

## Introduction

Understanding the functional structure of the nuclear pore complex (NPC) is not only of fundamental importance in biology but also invaluable for the design of polymer-based artificial nanopores(Jovanovic-Talisman et al., 2009; Hou et al., 2011; Tagliazucchi and Szleifer, 2015; Huang and Szleifer, 2017). As the largest cellular channel, NPC mediates the biomass transport between nucleus and cytoplasm with high selectivity and efficiency. Unlike mechanical or motor-driven biological nanochannels that undergo stimuli-responsive conformational transitions between open and closed states for gating, NPC has a relatively static scaffold constituted by the folded domains of hundreds of nucleoporins (Nups), and employs the intrinsically disordered regions (IDRs) of a subset of these Nups as its gatekeepers. Such IDRs form the central transporter(Beck, 2004; Kim et al., 2018; Yang et al., 1998), a selective permeability barrier that has been a long-standing black box due to the difficulty of experimental visualization(von Appen and Beck, 2016; Musser and Grunwald, 2016; Jovanovic-Talisman and Zilman, 2017). Termed as FG-Nups, the gating biopolymers use their phenylalanine-glycine (FG) repeat motifs to interact with nuclear transport receptors (NTRs) which facilitate the transport of macromolecules that carry specific labels (short peptides that serve as nuclear import/export signals)(Görlich and Kutay, 1999; Stewart, 2007; Köhler and Hurt, 2007). The self-associating propensity of FG-Nups, i.e., the hydrophobic pairing interaction between FG motifs, has led to the gel(Ribbeck and Gorlich, 2001; Frey and Görlich, 2007, 2009; Hülsmann et al., 2012; Schmidt and Görlich, 2016)-vs-brush(Hough et al., 2015; Lim et al., 2006; Rout et al., 2003, 2000; Wente and Rout, 2010) debate over the structure of the central transporter. While gelation of FG-Nups has been observed *in vitro*(Ader et al., 2010; Frey and Görlich, 2009), how these disordered biopolymers self-assemble *in vivo* under the geometrical constraints imposed by the scaffold is still an open question.

In the past few decades, several hypotheses of NPC gating have been proposed, such as virtual gate and polymer brush(Lim et al., 2006; Rout et al., 2000), selective phase(Ribbeck and Gorlich, 2001), reduction of dimensionality(Peters, 2005), and forest hypotheses(Yamada et al., 2010). As these hypotheses differ qualitatively in envisioning the functional structure of the central transporter that is currently unknown, a unified picture of NPC gating has not been established so far. It merits a note that all these hypotheses lack quantitative analysis of the spatial distribution of FG motifs and leave the electrostatic aspects of the central transporter unaddressed. In recent years, quantitative attempts(Ando et al., 2014; Gamini et al., 2014; Ghavami et al., 2016, 2014; Moussavi-Baygi and Mofrad, 2016; Osmanovic et al., 2013; Peyro et al., 2015; Tagliazucchi et al., 2013) have been made to elucidate the structure and functioning of the central transporter based on analytical models and molecular simulations. Prior work from our group, based on a molecular theory(Tagliazucchi et al., 2013) of the NPC, has shown that translocation across the NPC can be facilitated by NTRs that are both hydrophobic and negatively charged(Colwell et al., 2010). For the central channel, our previous model predicted a toroidal cloud of IDRs that has a higher density near the NPC scaffold than along the pore axis, in line with coarse-grained molecular dynamics (MD) simulations(Ghavami et al., 2014). Although the qualitative agreement between two different methods is encouraging, it is worth noting that the pore geometry and stoichiometry of the FG-Nups in previous models were based on early experimental findings that are now out of date. Recent experiments reveal that the NPC scaffold consists of three(Kim et al., 2018), rather than four rings(Alber et al., 2007), with the inner ring structure well preserved from yeast to human cells(Lin et al., 2016). For the yeast NPC, the stoichiometry (copy numbers of FG-Nups in the NPC) had nearly doubled in recent experimental reports(Kim et al., 2018; Rajoo et al., 2018) compared with previous ones(Alber et al., 2007). The implementation of such an experimental update is necessary for a faithful description of the central transporter by quantitative models.

In this paper, we revisit the functional structure of the central transporter of yeast NPC using a new molecular theory that has significant improvements in: 1) the geometry of the scaffold, 2) the stoichiometry and anchoring positions of the IDRs, 3) the description of molecular interactions, and 4) the differentiation between various FG motifs (see SI for details of the model). The new model explicitly considers the sequences of 11 FG-IDRs at the single-amino-acid resolution (Fig. 1A) and incorporates the latest experimental findings of the anchor positions (Fig. 1B). We classify the amino acids into 10 groups and model their hydrophobic, electrostatic, steric, van der Waals (vdW) interactions and acid-base equilibrium. We generate in total ∼10^8^ conformations of the IDRs to account for their conformational entropy. To examine the possibility of gelation of FG-Nups *in vivo*, we developed a method to account for the FG-FG pairing interaction, which was missing in our previous model. As schematically shown in Fig. 1A, we group all the FG motifs into three generic classes: 1) single FG motifs, 2) FG motifs with neighboring hydrophobic groups such as GLFG, and 3) FG motifs with separated hydrophobic groups such as FxFG. We also allow for attractions between specific spacers (amino-acid code N, Q, T) that were revealed by recent experiments(Ader et al., 2010) to be cohesive. In the Results and Discussions section, we show that the central channel of NPC incorporates inhomogeneous barrier structure, polarized electrostatic potential, well-separated charge-rich and charge-poor regions, and distinct domains of different FG motifs. Based on these insights, we discuss the sequence-structure-function relationship of the unfolded FG-Nups and propose a hypothesis of NPC-mediated transport that can be tested by future experiments.

**Figure 1.**
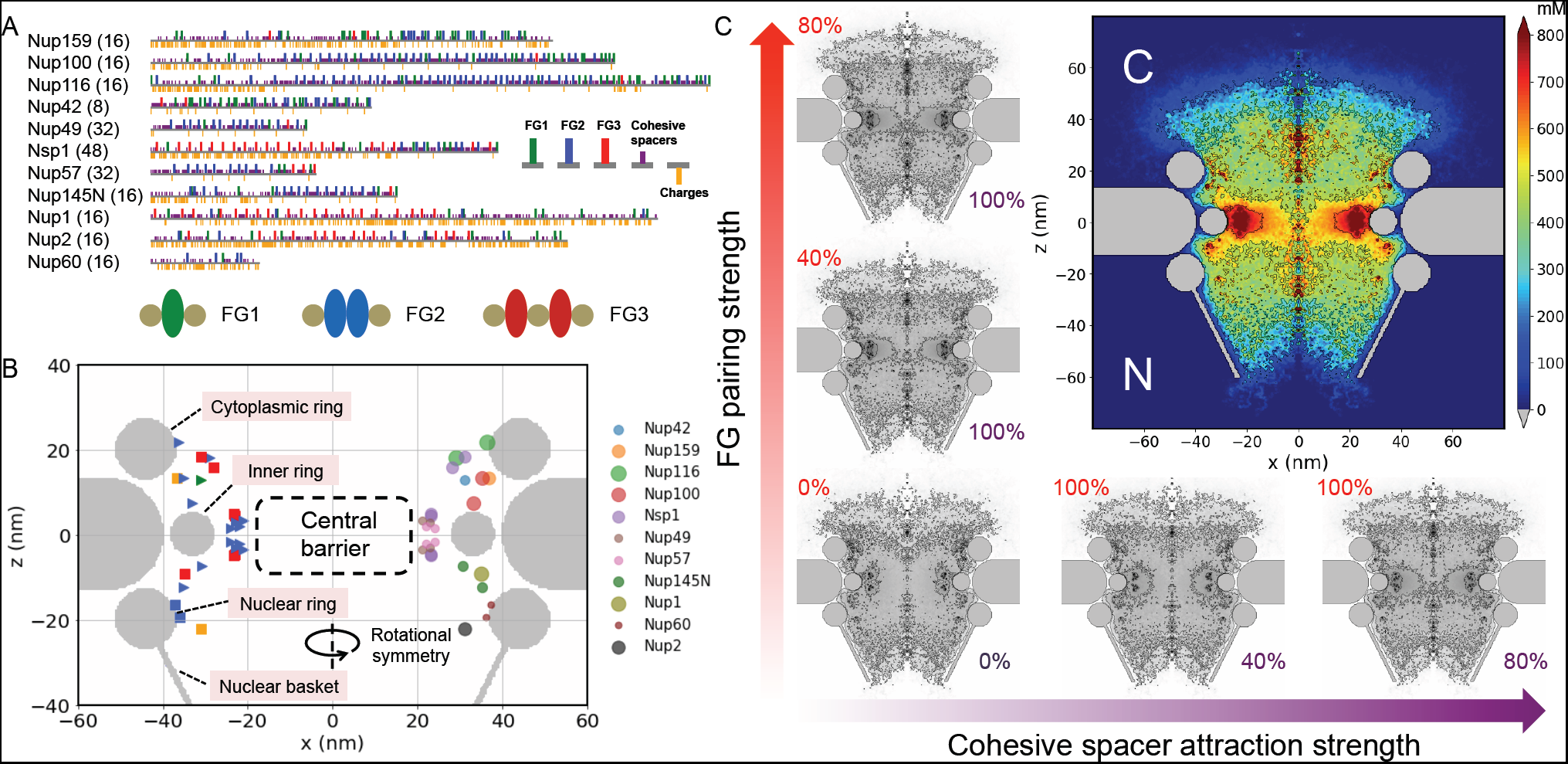
Basic input and output of the model. (**A**) IDR sequences with colored markers corresponding to different types of FG repeats and spacers. The names and stoichiometry of the FG-Nups are listed to the left of the anchoring ends of the sequences. Schematic representations of the tree types of FG motifs are shown under the sequences. Spheres indicate hydrophilic and ovals hydrophobic amino acids. (**B**) Geometry of the model NPC. The scaffold rings are coarse-grained into three tori. On the right, the anchor positions of the IDRs are represented by colored discs (disc size indicates the length of the sequence). On the left, the color of the anchor position aligns with the color code of the dominating FG-type of the IDR (colored orange if there is no dominant FG-type). Triangles indicate cohesive IDRs, squares non-cohesive and Nsp1. (**C**) Color map (upper right) of the mM concentration of all the amino acids inside the NPC with cooperative FG-pairing (2.5kT) and cohesive spacer attraction (1kT). Structural dependence of the central channel on FG-paring and spacer attraction strengths is shown in the grey panels with the normalized FG-pairing strength and cohesive spacer attraction strength displayed in the upper left and bottom right corners, respectively.

## Results and Discussions

### Cooperation of FG-pairing and specific spacer attraction in shaping the central transporter

Experiments suggest that the FG-pairing interaction and the attraction between the cohesive (NQT) spacers are two major driving forces for self-association of FG-Nups(Patel et al., 2007). It has been also observed, as clearly demonstrated by the color-coded sequences in Fig. 1A, that the cohesive domains of Nup116, Nup100, Nup145N, Nup57, and Nup49 contain mostly NQT amino acids and type-2 FG motifs. Despite the lack of quantitative data for the energetics of the FG-pairs, both NMR measurements(Hough et al., 2015) and all-atom MD simulations(Raveh et al., 2016) suggest that interaction between FG motifs is weak and highly dynamic, consistent with their hydrophobic nature. We have carried out MD simulation to show that Phe-Phe pairing energy (between single-molecule amino acids) is around 2.5kT in water at 300K (Fig. S1). Besides FG-pairing, we also assigned 1kT interaction energy between the cohesive NQT spacers (see SI for more details). Before we focus on the reasonably cohesive conditions, it is instructive to systematically study the molecular organization of FG-Nups under various arbitrary combinations of the FG-pairing and spacer cohesiveness. As shown in the lower left panel of Fig. 1C, when all the cohesive interactions are turned off, the overall spatial distribution of the IDRs is highly diffuse with the density of amino acids being lower along the pore axis than near the scaffold where the IDRs are anchored. Increasing the FG-pairing strength and the spacer cohesiveness contracts the FG-Nups into the central barrier zone encircled by the inner scaffold ring (location marked in Fig. 1B), where gelation is expected to happen according to the selective phase hypothesis(Hülsmann et al., 2012). Notably, the predicted condensation is rather limited if one of the two cohesive forces is weak, in line with *in vitro* experimental observations that both FG-pairing and attractive spacer interaction are indispensable for enabling gel-like barrier structures(Ader et al., 2010; Patel et al., 2007; Xu and Powers, 2013). However, we found that even with both relatively strong FG-pairing (2.5kT) and spacer attraction (1kT), i.e., condition for the color panel in Fig. 1C, the central barrier does not seal itself and leaves open a narrow axial conduit. Such unoccluded barrier structure near the inner ring is similar to that found in electron microscopy (EM) experiments(Eibauer et al., 2015), and is consistent with the single-molecule super-resolution fluorescence observation of a single central channel for passive diffusion of small molecules(Ma et al., 2012). Our model predicts that the morphology of the central barrier is sensitive to the nature of the cohesive interaction: strong FG-pairing tends to homogenize the IDR spatial distribution through network formation whereas specific spacer attraction tends to collapse the barrier into high-density condensates, rendering a heterogeneous gating structure (Fig. S5). While in our model the interactions between FG-Nups are determined by their intrinsic chemical properties, they can be effectively modulated by NTRs through multivalent binding and electrostatic interactions. It has been observed in experiments that NTRs can induce morphological alternation of the assemblies of FG-Nups(Lim et al., 2007; Wagner et al., 2015). Fluorescence experiments reported(Ma et al., 2012) that the central passive pathway widens upon raising the concentration of Importin-*β* (Imp*β*), a primary NTR responsible for nuclear import.

Outside the central barrier, our model (with both FG pairing and cohesive spacers) predicts two lobe-shaped high-density regions, reminiscent of recent EM studies where the central transporter appears as a two-lobed blur(Kim et al., 2018). The condensation of FG-Nups at the two exits of the pore implies that the functional gate of NPC is not limited to the central barrier but extend to the cytoplasmic and the nuclear sides. In particular, the prediction of a prominent cohesive zone at the cytoplasmic vestibule of the NPC suggests that molecular screening for nuclear import may take place before the cargoes reach the central barrier of NPC. Apart from the condensed zones, the overall spatial distribution of FG-Nups is diffuse enough to create a highly dynamic FG cloud encapsulating the central barrier, in accord with the atomic force microscopy (AFM) observations(Sakiyama et al., 2016) of large structural variance of the FG-Nups looking from the cytoplasmic side of NPC. It is worth noting that since the vestibular condensation of FG-Nups at the cytoplasmic exit of the pore requires strong cohesiveness to compensate for the conformational entropy penalty, such condensation may not be feasible without the aid of Imp*β*. In line with the above consideration, a recent AFM study reported that Imp*β* facilitates the occlusion of the cytoplasmic side of NPC(Stanley et al., 2018). Note also that while the long cohesive FG-Nups (Nup116, Nup100) anchored at the cytoplasmic side can extend into the NPC and seal the central barrier in conjunction with short cohesive FG-Nups (Nup57, Nup49) that emanate from the inner ring, this would involve a conformational entropy penalty and is therefore unlikely, as indicated by our model in which both energy and conformational entropy considerations are quantitatively taken into account. The short FG-Nups alone cannot occlude the entire central barrier zone due to the geometrical constraints, as demonstrated by experiments on artificial nanopores that mimic the NPC(Kowalczyk et al., 2011). Nevertheless, despite the unsealed morphology of the central barrier, its ring structure is more condensed than what would be expected for a non-cohesive polymer brush envisioned by the virtual gate model.

### Thermoreversible FG-network and asymmetric electrostatic potential

The crux of the brush-gel debate lies in the degree of crosslinking of the FG-Nups, or the pairing fraction of FG-repeats(Hülsmann et al., 2012), which holds the key to the functional structure of the central transporter. In Fig. 1C (color pannel) we have shown how heterogeneous barrier structures emerge under cooperative FG (2.5kT) and spacer (1kT) cohesiveness. In the remainder of the paper, we focus on this reasonably cohesive case and visualize its fine structure from a diversity of perspectives that the model provides, starting with the spatial distributions of FG repeats. As shown in Fig. 2A, our model predicts a diffuse yet inhomogeneous spatial distribution of FG-repeats. The FG concentration reaches around 40 mM inside the condensed domains and drops to 10-30 mM outside them. The overall FG concentration is lower than the ∼50 mM saturation limit suggested by *in vitro* experiments(Frey and Görlich, 2007), but is significantly higher than the estimates (<10 mM) from super-resolution fluorescence experiment(Ma et al., 2016). Even for the non-cohesive system that features a brush-like morphology, we found an average FG concentration in the range of 20-30 mM (Fig. S3B). If the current estimates of the stoichiometry of FG-Nups are reliable(Kim et al., 2018; Rajoo et al., 2018), the average concentration cannot be much lower than that, due to the confinement imposed by the scaffold. This means that the average FG concentration is not sensitive to whether the morphology of the central transporter is brush-like or gel-like. The quantity that truly distinguishes the two cases is the FG-pairing fraction, which depends not only on the FG concentration but also on the FG interaction strength, and can go from nearly none (completely non-cohesive) to almost 100% (saturated pairing).

**Figure 2.**
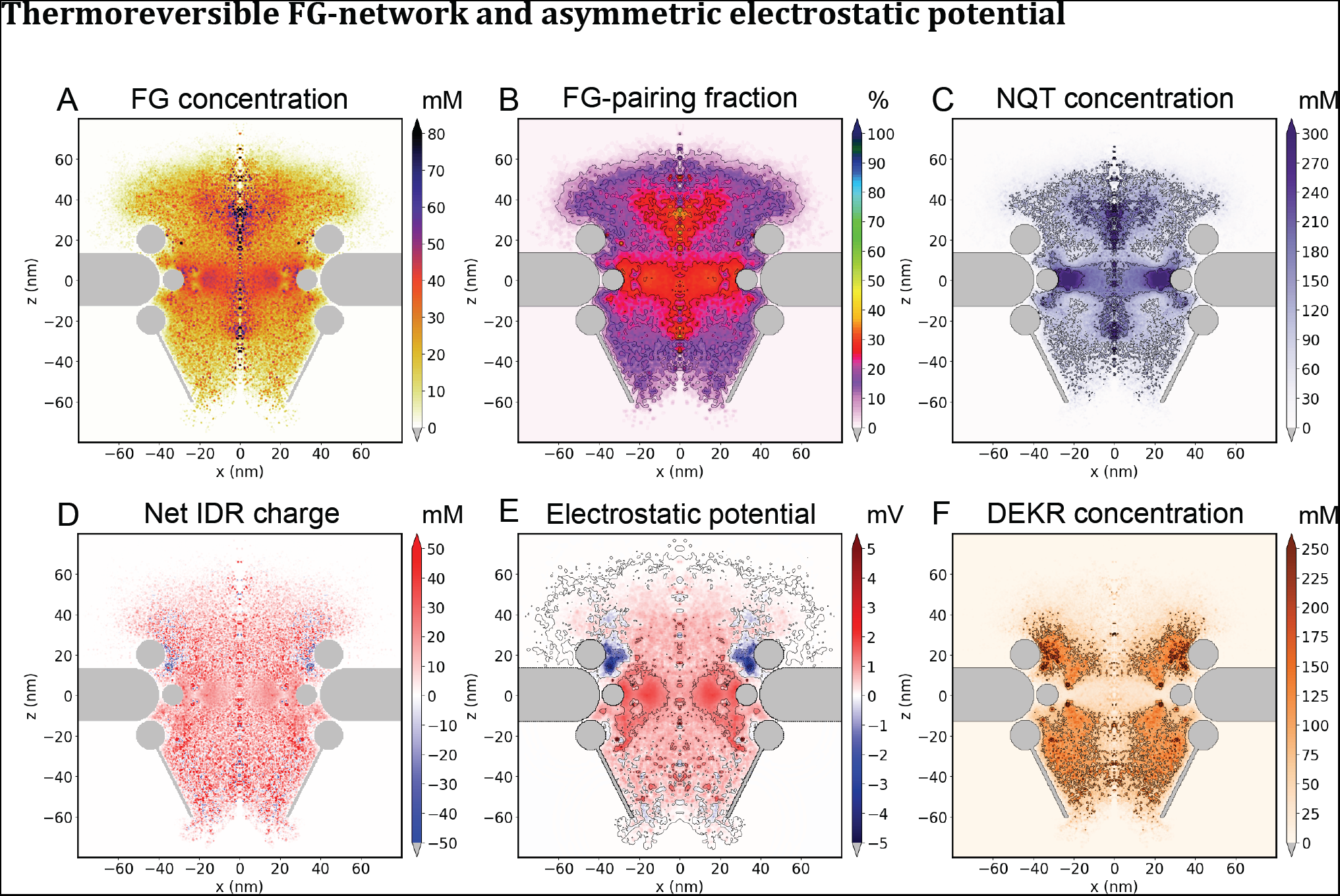
Central transporter as a FG-rich and charge-poor region. (**A**) Overall FG concentration throughout the NPC in mM. (**B**) FG-pairing fraction in percentage. (**C**) Concentration of NQT (cohesive) spacers in mM. (**D**) Concentration of net IDR charge in mM. (**E**) Electrostatic potential throughout the NPC in mV. (**F**) Concentration of DEKR (charged) spacers in mM.

Fig. 2B shows our theoretical predictions for the FG-pairing fraction throughout the NPC. On average the pairing fraction is around 30%, which is an order of magnitude higher than that obtained for a non-cohesive system (∼3%, Fig. S3C). This number, however, is well below the saturation limit assumed in the selective phase model(Frey and Görlich, 2007), which means there are many (∼70%) dangling FG motifs that are ready to bind with NTRs. In a sense, the thermoreversible FG-network predicted by our model is in an intermediate state between a brush and a gel, with both brush and gel characteristics to some degree. The pairing fraction is not homogeneous and exhibits a spatial pattern that overlaps with the FG-rich domains in Fig. 2A, reflecting the fact that FG-pairing tends to condense FG motifs. Moreover, regions rich in FG motifs and high pairing fraction roughly coincide with domains rich in cohesive NQT spacers (Fig. 2C), highlighting again the important role of these spacers in shaping the central transporter. Figs. 2A-C reveal the existence of two vestibules at both the cytoplasmic and the nuclear sides that are rich in FG motifs and cohesive spacers, which could recruit Imp*β*1 at both exits of the central pore, as observed in experiments(Lowe et al., 2010). In our recent theoretical study of the transport pathways of model cargoes through a cylindrical nanochannel coated with homopolymers, we found that the cargoes with moderate polymer affinity tend to accumulate near the two vestibules of the channel, due to the substantially larger accessible volume (and therefore larger entropy) for both the cargoes and the polymers(Tagliazucchi et al., 2018). The shape of the NPC scaffold with widely open exits and the deployment of long FG-Nups at the outer rings suggest that a pooling mechanism (vestibular accumulation for efficient transport) has been exploited and optimized in the nucleocytoplasmic transport. Such pooling of NTRs could in return strengthen the vestibular barriers to block unrecognized macromolecules.

In addition to the thermoreversible FG-network, we predict a net positive charge homogeneously distributed throughout most of the central transporter (except near the cytoplasmic ring, see Fig. 2D), with an average net charge concentration of about 20 mM. Such positively charged nano-environment is electrostatically favorable for macromolecules that are negatively charged, consistent with the prior finding that NTRs and NTR-cargo complexes bear more negative charges than most cellular proteins(Colwell et al., 2010). To better understand the electrostatics of the NPC, we calculated the electrostatic potential produced by the charged FG-Nups. As shown in Fig. 2E, the overall potential is positive as expected based on the net charge distribution. However, it is intriguing that this self-built potential is highly inhomogeneous and asymmetric in space, in a way that is not correlated with the spatial heterogeneity of FG-repeats, revealing another dimension of the molecular organization inside the NPC. Compared to the relatively weak and uniform potential in the pore center, the potential near the scaffold is both intensified and polarized. In particular, a negative potential appears near the cytoplasmic ring and transitions into positive potential near the inner ring and the nuclear ring. The roughly 1 mV difference between the inner scaffold ring and the axis of the pore is expected to provide an electrostatic energy bonus of about 2kT for the NTRs of average charge around −50e, to follow a peripheral pathway near the inner ring. This could explain fluorescence and EM observations of NTRs such as Imp*β*1, NTF2, Kap104 and Kap121 near the periphery of the pore(Fiserova et al., 2010; Ma and Yang, 2010; Ma et al., 2012), and fluorescence observations that positively charged cargoes pass the NPC along the axial channel(Ma et al., 2016). Note that the observation of peripheral translocation of NTRs is difficult to explain from a FG-binding perspective alone, given the relatively homogeneous distribution of FG motifs in the central barrier (Fig. 2A). Our prediction highlights the possible role of the self-consistent electrostatic field in tuning the NTR pathway, a mechanism that has not been addressed by previous models. We propose that the center-to-periphery electrostatic potential gradient participates in dispersing cargoes according to their charge to size ratios. The functional role of the negative potential near the cytoplasmic ring is not entirely clear at present but it is likely to assist with NTR pooling before nuclear import and to direct the negatively charged cargoes to the central ring. It is worth noting that the polarized electrostatic potential arises not only due to the net charge distribution but also due to the inhomogeneous osmotic pressure inside NPC. In fact, in stark contrast to the net charge distribution, the distribution of charged amino acids (DEKR, positive and negative, see Figs. S2A, B) of the IDRs has a highly inhomogeneous spatial pattern (Fig. 2F), in remarkable anti-correlation with the neutral cohesive spacers (Fig. 2C), suggesting phase separation between the charge-rich non-cohesive and the charge-poor cohesive regions inside the nuclear pore.

### Spatial segregation of different FG motifs and an atlas of individual FG-Nups

Figs. 2A, B depict a thermoreversible FG-network where unpaired FG-motifs are widely dispersed and available for binding of NTRs throughout the NPC. However, at this point it is still unclear how such a diffuse cloud of FG-motifs directs the traffic through the lumen of the NPC. To shed more light on this issue, we distinguish between three generic types of FG motifs. Type-1 contains one phenylalanine as the only hydrophobic motif, mostly of FG-type. Type-2 has neighboring hydrophobic groups such as GLFG, xAFG and, xIFG, whereas type-3 has separated hydrophobic groups such as FxFG, LSFG, ISFG (x indicates neutral hydrophilic amino acids only, since neighboring charged amino acids are expected to suppress hydrophobicity(Huang et al., 2015)) The spatial distributions of the three types of FG motifs are shown in Fig. 3A-C. It is interesting that the different FG motifs form distinct nano-domains in space. The single FG motifs are concentrated along the axis of the pore (Fig. 3A), filling the low-density axial conduit we showed in Fig. 1C, which could explain the experimental finding that the central channel for the passive diffusion of small molecules is more viscous than an open aqueous conduit(Ma et al., 2012). The central and vestibular barriers are enriched predominantly by type-2 FG motifs (Fig. 3B), whereas most type-3 FG motifs (Fig. 3C) are widely distributed outside the barriers. Figs. 3E, F show the spatial distributions of GLFG, FxFG, the most-studied type-2 and type-3 FG motifs, which are clearly segregated from each other. The spatial distribution of other FG motifs (non-GLFG-FxFG) is peaked about axis of the NPC, similarly to the single FG motifs (Fig. 3D). The complementary nano-domains of distinct FG motifs are expected to add on the electrostatic potential another layer of pathway selectivity and specificity(Curk et al., 2017) for multivalent NTRs and their cargo-complexes to undergo path-selective transport.

**Figure 3.**
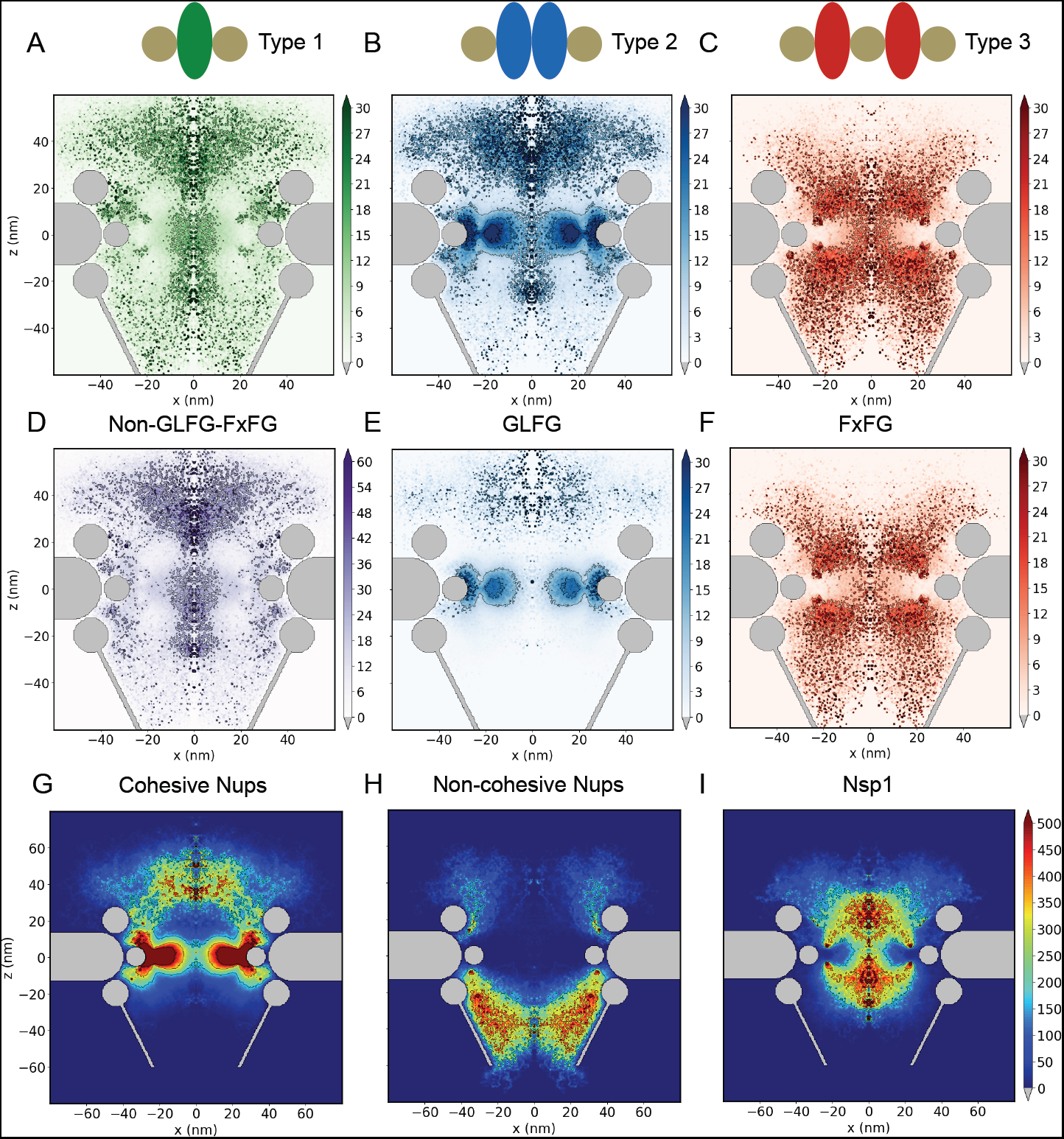
Distinct domains of different FG motifs and of cohesive and non-cohesive FG-Nups. (**A-C**) Spatial distributions of three generic types of FG motifs (see main text for the classification protocol). (**D-F**) Spatial distributions of non-GLFG-FxFG (including type-1) motifs, GLFG (belonging to type-2) motifs, and FxFG (belonging to type-3) motifs. Note that panel D has different concentration scale than panels E, (**G-I**) Spatial distributions of cohesive FG-Nups (Nup116, Nup100, Nup42, Nup57, Nup49, Nup145N), non-cohesive FG-Nups (Nup159, Nup1, Nup2, Nup60) and Nsp1 (partially cohesive). All the color maps show amino-acid concentrations in mM.

It is worth noting that all FG motifs have similar pairing energy in our model. Therefore, the segregation of different FG motifs is not a trivial liquid-liquid phase separation but is programmed into the amino-acid sequences of the FG-Nups. Under our FG classification protocol, subdomains of type-2 and type-3 FG-motifs can be clearly seen in the color-coded sequences shown in Fig. 1A. Moreover, the two types of subdomains have distinct concentrations of cohesive spacers (purple) and charges (orange). It is well known from *in vitro* experiments that GLFG-rich Nups such as Nup116, Nup100, Nup57, Nup49 contain most cohesive subdomains(Patel et al., 2007) that are vital for forming the permeability barrier. Recent experiments reveal that GLFG-motifs directly bind to multiple scaffold Nups and that the GLFG-rich long Nup116 and Nup100 play important roles in the biogenesis of the NPC(Onischenko et al., 2017). In our model, we have assigned weak interactions between the inner surface of the coarse-grained scaffold and all the FG-Nups. In line with the experimental observations, we predict that GLFG-rich Nups are localized in the vicinity of the scaffold and constitute the cohesive central barrier (Fig. 3G). Remarkably, our model predicts that long Nup116 and Nup100 form a cytoplasm-oriented sieve-like structure, analogous to the nuclear basket but more disordered. The overall spatial distribution of the cohesive FG-Nups is also cytoplasm-oriented, suggestive of a potential role of the spatial gradient of type-2 FG-motifs in guiding nuclear export. Interestingly, it has been observed by super-resolution imaging that Nup116 segment as a cargo(Ma et al., 2016) (which can homotypically interact with GLFG-Nups) and mRNA during export(Ma et al., 2013) both have a similar spatial pattern with high dwelling probability in the central barrier ring and the cytoplasmic vestibule. On the other hand, the larger amount of type-3 FG motifs within the nuclear half of the NPC suggests that their spatial gradient could direct nuclear import, in line with reports that FxFG motifs are stronger binders to the hydrophobic pockets of Imp*β* than GLFG motifs(Isgro and Schulten, 2005). Fig. 3H presents the spatial distribution of non-cohesive FG-Nups, which shows up in the periphery of the cytoplasmic half and fills the nuclear half of NPC. The partially cohesive Nsp1 with non-cohesive FxFG-rich subdomain near the anchoring end and cohesive subdomain near the free end, fills the central lumen of the NPC, while depleted from the scaffold and the central barrier (Fig. 3I).

Fig. 4 shows an atlas of 11 types of individual FG-Nups. The cytoplasm-oriented, center-oriented and nucleoplasm-oriented FG-Nups are displayed in the upper, middle and lower rows, respectively. The spatial distributions of the FG-Nups along the axis of the pore are largely determined by their anchor positions. The central FG-Nups have more copy numbers than the cytoplasmic and nuclear ones. Among them, the Nup49 and Nup57 are short in length and constitute the high-density central ring rich in GLFG motifs. On the cytoplasmic side, Nup116 and Nup100 participate in forming the cytoplasmic barrier of the pore whereas Nup159 and Nup42 reside at the pore periphery, consistent with the experimental observation that Nup116 and Nup100 contribute more to the NPC permeability barrier than other FG-Nups(Timney et al., 2016). Note that while Nup116 has both swollen and collapsed subdomains, the collapse of its cohesive subdomain tends to happen near the pore axis. Nup159 carries more negative charges than positive ones and contributes to the negative electrostatic potential shown in Fig. 2E. It is interesting to observe how these highly charged long FG-Nups extend into the cytoplasmic side like antennas. In a non-cohesive system (Fig. S3), Nup116 and Nup100 do not block the cytoplasmic side and have a peripheral distribution like that of Nup159. Near the nucleoplasmic side, the FG-Nups also differ in their lengths and spatial distributions. The long IDRs of Nup1 and Nup2 are enriched in type-3 FG motifs, in contrast to the short IDRs of Nup145N, Nup60 that carry mostly type-2 FG-motifs. It is worth noting that, except for the most abundant Nsp1, all the FG-Nups have localized spatial distributions and are characterized by specific FG motifs. The lack of overlap between the cytoplasm- and nucleoplasm-oriented FG-Nups suggests that nucleocytoplasmic transport necessitates switching between different FG-Nups by a sequence of binding and unbinding events.

**Figure 4.**
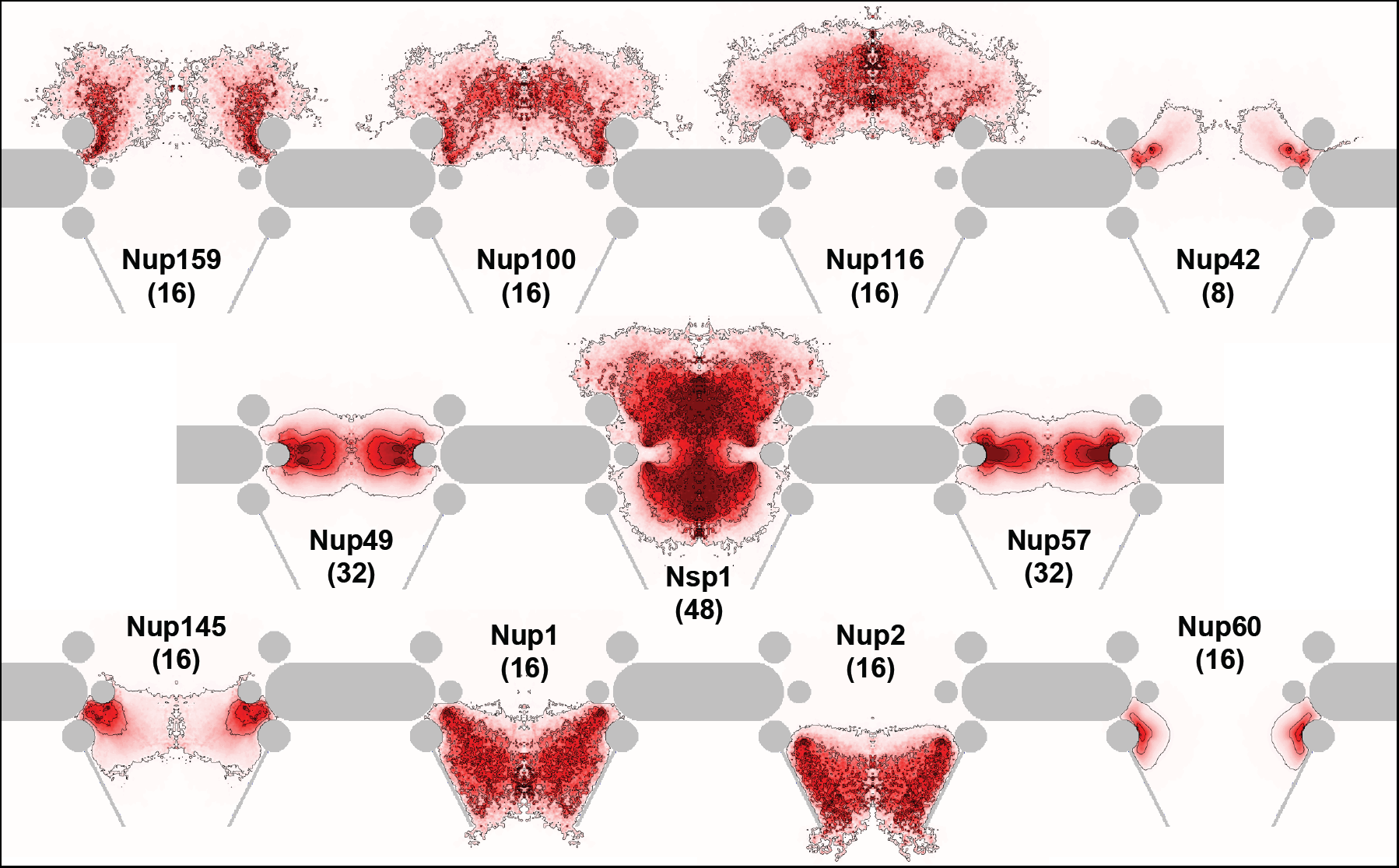
An atlas of various FG-Nups of yeast NPC shown in color maps. From top to bottom, the three rows show the spatial distributions of the FG-Nups with their anchor positions located towards the cytoplasm, near the central inner ring, and towards the nucleoplasm. The copy number of each individual Nup is indicated in the parenthesis.

### The whole is more than the sum of its parts

The current advances in revealing the structure of the NPC scaffold have been based on a divide-and-conquer methodology which breaks this structure into subcomplexes that can be analyzed at atomic resolution using protein crystallization and then integrated back to get the whole picture. Can we apply an analogous approach to understand the functional core, i.e., the central transporter of the NPC?

To answer this question, we studied a reference system where isolated IDRs are characterized individually and superposed to construct an overall gating structure. In other words, the cross-interactions between different IDRs are turned off in this reference system. Fig. 5A shows the overall gating structure of the reference system and a few typical spatial distributions of the isolated IDRs. Since most of the FG-Nups have anchoring positions within 20nm of the pore equator (Fig. 1B), the reference system has a concentrated IDR distribution inside the central barrier zone and near the inner ring of the scaffold. However, compared to the fully interacting system (Fig. 1C, color panel), it lacks the vestibular condensates at the exits of the pore, suggesting that the formation of vestibular barriers/recruiters necessitates the interplay between different FG-Nups, and especially the volume exclusion between different IDR territories (Fig. 4). In the fully interacting system, the spatial distributions of long FG-Nups such as Nup100, Nup116, Nup159, Nup1 and Nup2 are extended towards either the cytoplasmic or the nuclear side of the NPC depending on their anchoring positions (Fig. 3G, H, Fig. 4), whereas Nsp1 with anchoring positions across the pore equator have polarized distributions that are depleted around the central barrier ring (Fig. 3I, Fig. 4). In the reference system, these FG-Nups in their isolated states tend to occupy the NPC lumen in a less segregated way (Fig. 5A). Among all the isolated FG-Nups, Nup116 are predicted to form the largest condensate along the pore axis (Fig. 5A), consistent with their leading role in NPC biogenesis. Compared to the fully interacting system (Fig. 2), the reference system has drastically different spatial distributions of the cohesive (NQT) and charged (DEKR) spacers (Fig. 5B, C), with no sign of phase separation between them. The net charge of FG-Nups is less homogeneously distributed (Fig. 5D) and the electrostatic potential is more intensified in the central barrier (Fig. 5E). The reference system has a more concentrated distribution of all the FG motifs and has more intermixed domains of distinct FG motifs (Fig. 5F-I), compared to the fully interacting system (Fig. 2A, Fig. 3A-C). These comparisons highlight the importance of cross-interaction between FG-Nups in forming the extensive and intricate gating structure of NPC, which demonstrates that the central transporter as a whole is more than the sum of the parts. Consistent with deletion experiments(Strawn et al., 2004), our result suggests that deleting a sufficiently large number of FG-Nups in the pore will affect the overall function, even if those FG-Nups are not directly involved in the transport mechanism for a given NTR.

**Figure 5.**
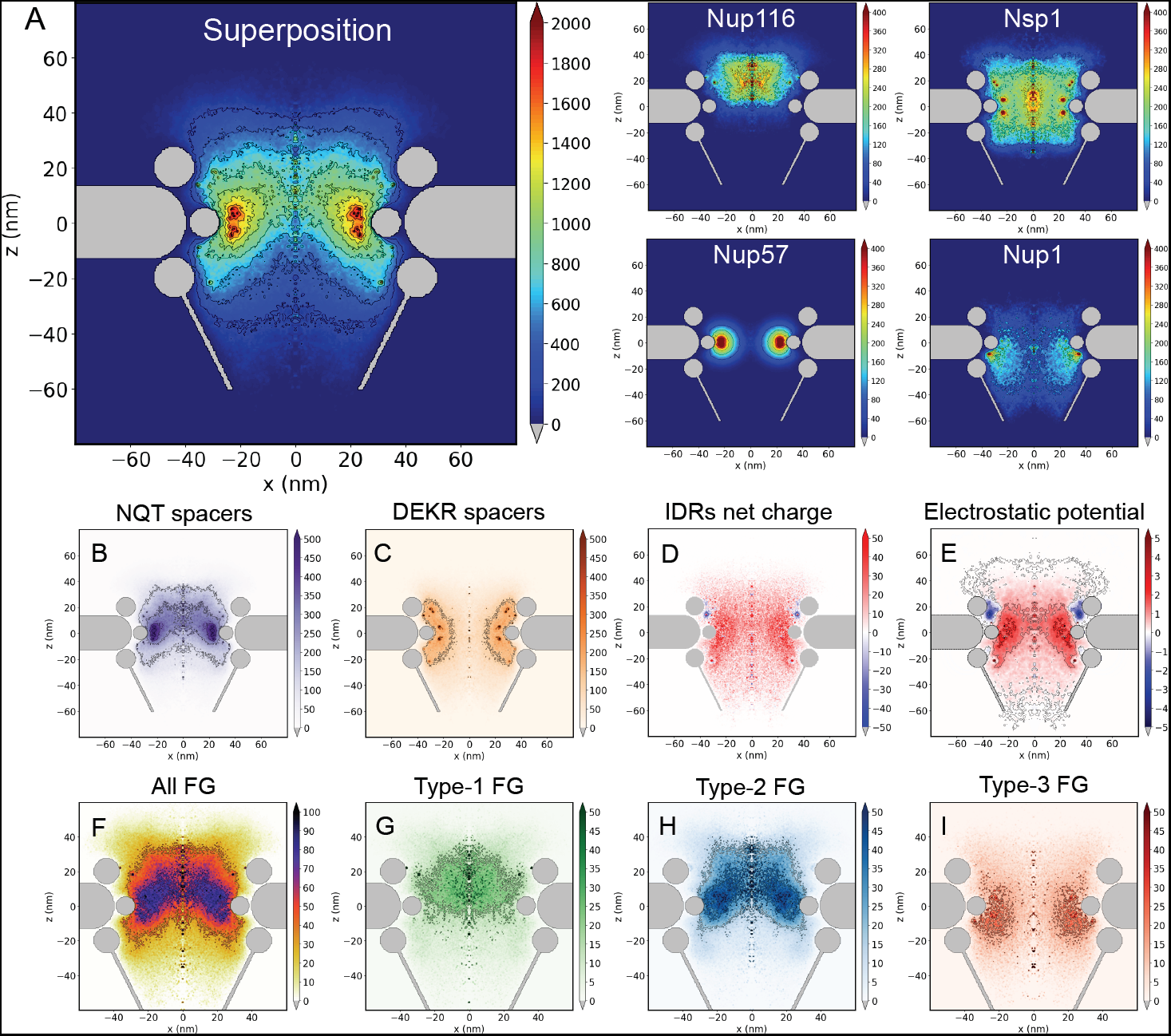
Reference system of non-interacting FG-Nups. (**A**) Superposition of all 11 IDRs (large panel) and 4 typical individual IDRs (small panels). (**B-I**) Spatial distributions of NQT spacers (B), charged DEKR spacers (C), net charge of IDR amino acids (D), electrostatic potential (E), all FG motifs (F), type-1 FG motifs (G), type-2 FG motifs (H) and type-3 FG motifs.

## Conclusion and outlook

In this work, we have studied a molecular model that provides high-resolution structural details about the distribution of intrinsically disordered regions (IDRs) inside the NPC. Our results reveal an intricate integration of various FG-Nups, resulting in an elaborate central transporter. Besides a high-density FG-ring at the equator of the pore that has been reported in previous models(Ghavami et al., 2014; Tagliazucchi et al., 2013), our work suggests the existence of vestibular condensates along the axis of pore that can serve as barriers for inert molecules and as attractors for FG binders. Our quantitative analysis suggests that the permeability barrier extends from near the inner ring of the scaffold to the two exits of the NPC, forming a 3dz^2^-orbital-shaped structure. The vestibular condensation of FG-Nups is in accord with the experimentally observed pooling of NTRs(Lowe et al., 2010), which could in return strengthen the distal barriers at the pore exits.

Our analysis highlights the importance of the cohesive domains laden with specific attractive spacers (NQT) in guiding the self-assembly of FG-Nups *in vivo* into segregated cohesive and non-cohesive zones with the latter being rich in charges. However, even inside the cohesive region we find the pairing fraction of FG motifs to be less than 50%, meaning there exist more dangling than paired FG motifs. In concord with recent experimental finding(Patel et al., 2007), we predict that FG-Nups that are rich in cohesive subdomains such as Nup116, Nup100, Nup57, Nup49 and Nup145N, dominate the proximity of the NPC scaffold and are crucial to the permeability barrier. Our model further suggests that Nup116 and Nup100 could form a sieve-like structure towards the cytoplasm, consistent with their important roles in NPC biogenesis(Onischenko et al., 2017). However, the overall spatial distribution of the FG-Nups is predicted to remain diffuse in the cytoplasmic side, meaning the cytoplasmic part of the NPC is highly flexible and dynamic, in line with AFM observations(Sakiyama et al., 2016). It is possible that NTRs are needed to elicit or stabilize the predicted cytoplasmic-sieve structure(Stanley et al., 2018). By classifying the FG motifs into three generic groups, we find the cohesive subdomains to be rich in type-2 FG motifs with neighboring hydrophobic amino acids, such as GLFG, xAFG, xIFG. While it is well known that GLFG are crucial for the cohesiveness of FG-Nups, more experimental efforts are needed to investigate whether other type-2 FG-motifs facilitate barrier formation and NPC biogenesis.

Our model reveals an intensified and polarized electrostatic field near the NPC scaffold. The highly positive potential near the inner ring provides an electrostatic explanation for the experimental finding that negatively charged NTRs tend to shuttle near the NPC scaffold(Fiserova et al., 2010; Ma and Yang, 2010; Ma et al., 2012) whereas positive cargoes are confined to the axial channel(Ma et al., 2016). On the other hand, the unoccluded central barrier predicted by our model is consistent with the experimental observation that passive diffusion of small cargoes takes an axial pathway(Ma et al., 2012). The unsaturated FG-network predicted by our model features complementary nano-domains of different FG motifs, implying their distinct functions in the selective barrier. The compartmentalization of FG motifs is encoded in the amino-acid sequences of the IDRs and in the anchoring addresses at which they emanate from the scaffold and does not incur significant conformational entropy penalty for the FG-Nups. However, we show that interactions between different FG-Nups are necessary to orchestrate and sustain such organized FG-territories.

We propose that the combination of multivalent targeting, hydrophobic and electrostatic interactions and entropic effects allows the central transporter of NPC to control the pathways of cargoes according to their FG-affinity, charge, and size. Along their peripheral pathway favored by the electrostatic interaction, the NTRs will likely need to transition between the FxFG and GLFG domains, with small energetic gain or loss through multivalent weak hydrophobic interactions. Such multivalent targeting scenario has been recently shown to enable high molecular sensitivity and specificity compared to strong monomeric binding(Curk et al., 2017). For NTRs that have a higher affinity to FxFG than to GLFG, for example Imp*β* as suggested by literature(Isgro and Schulten, 2005), passing through the GLFG ring will have counteracting energetic effects from FG-binding and electrostatic interaction that permit fast trafficking, whereas NTRs that are more GLFG-philic could be trapped near the scaffold. Besides FG-binding and electrostatics, the size of the cargo is another factor that influences the path-selective transport. In our recent theoretical study, we found that entropic effects drive large cargoes to take a more centralized pathway through a polymer-coated nanochannel, and to pool at the channel exits due to cargo-polymer affinity(Tagliazucchi et al., 2018). The pooling mechanism could accelerate the tunneling of large cargoes through NPC. More systematic experimental investigation on the specific NTR-FG interactions is needed towards a full picture of path selectivity.

In summary, our model pivots a wide array of existing experimental observations, and reconciles the conundrum between high efficiency and high specificity of nucleocytoplasmic transport by predicting: 1) a diffuse thermoreversible (weakly and partially cross-linked) FG-network with widely available dangling FG motifs for fast NTR binding and unbinding, 2) complementary nano-territories of distinct FG motifs and an inhomogeneous electrostatic potential that can cooperate to direct the transport pathway through the combination of multivalent FG-targeting and electrostatic steering. These results shed light on the sequence-structure-function relationship of the unfolded FG-Nups, which can be tested by new experiments. Future modeling efforts will be directed towards the study of transport dynamics through the predicted NPC structure.

## Acknowledgements

IS, KH gratefully acknowledge funding from NSF Biol & Envir Inter of Nano Mat 1833214, and NIH National Cancer Institute R01 CA228272. YR would like to acknowledge support by grants from the Israel Science Foundation and from the Israeli Centers for Research Excellence program of the Planning and Budgeting Committee. We thank Dr. André Hoelz and Dr. Barak Raveh for discussion of the NPC scaffold.

## Author contributions

Conceptualization, K.H., Y.R., and I.S.; Methodology, K.H.; Software, K.H., M.T., and S.H.P.; Formal Analysis, K.H.; Investigation, K.H.; Writing-Original Draft, K.H.; Writing-Review & Editing, K.H., Y.R., M.T., S.H.P., and I.S.; Supervision, Y.R., and I.S.; Project Administration, I.S.; Funding Acquisition, Y.R., and I.S.

## Declaration of interests

The authors declare no competing interests.

