## Supplemental Information for "Molecular model of the nuclear pore complex reveals a thermoreversible FG-network with distinct territories occupied by different FG motifs"

### Molecular theory

To model the nuclear pore complex (NPC), we use a molecular theory that incorporates the intrinsically disordered regions (IDRs) sequences of the FG-Nups at the amino-acid resolution and explicitly accounts for the conformational, amphiphilic, electrostatic, steric and acid-base properties of the IDRs. The molecular theory combines both field and molecular representations of the system and solves the free energy functional using a self-consistent numerical approach. The free energy functional purposed for the central transporter of the NPC is written as follows:

$$
\begin{aligned}
\beta F = & \sum_{i\in \mathrm{A,C,w}} \int d\mathbf{r}\rho_i(\mathbf{r})\left[\ln\left(\rho_i(\mathbf{r})v_{\mathrm{w}}\right)-1\right] + \sum_{i\in \mathrm{H^+,OH^-}} \int d\mathbf{r}\rho_i(\mathbf{r})\left[\ln\left(\rho_i(\mathbf{r})v_{\mathrm{w}}\right)-1+\mu_i^0\right] \\
& + \int d\mathbf{s} \sum_{k=1,N_k} \sigma_k\left(\mathbf{r}(s)\right)\sum_{\alpha} P_k\left(\mathbf{r}(\mathbf{s}),\alpha\right)\ln\left(P_k\left(\mathbf{r}(\mathbf{s}),\alpha\right)\right) \\
& + \sum_{l=1,N_l}\sum_{l'=1,N_l} \frac{\beta\varepsilon_{ll'}}{2}\iint d\mathbf{r}\,d\mathbf{r}'\,g\left(\left|\mathbf{r}-\mathbf{r}'\right|\right)\left\langle n_l(\mathbf{r})\right\rangle\left\langle n_{l'}(\mathbf{r}')\right\rangle \\
& + \int d\mathbf{r}\left[\left\langle \rho_Q(\mathbf{r},t)\right\rangle \beta\psi(\mathbf{r}) - \frac{1}{2}\varepsilon\left(\nabla\psi(\mathbf{r})\right)^2\right] \\
& + \sum_{l=1,N_l}\int d\mathbf{r}\left\langle n_l(\mathbf{r})\right\rangle\left[f_l(\mathbf{r})\left(\ln\left(f_l(\mathbf{r})\right)+\beta\mu^0_{l,\mathrm{charged}}\right)+\left(1-f_l(\mathbf{r})\right)\left(\ln\left(1-f_l(\mathbf{r})\right)+\beta\mu^0_{l,\mathrm{uncharged}}\right)\right] \\
& + \sum_{p\in \mathrm{FG,NQT}}\int d\mathbf{r}\left\langle n_p(\mathbf{r})\right\rangle\left[\frac{g_{\mathrm{p}}(\mathbf{r})}{2}\ln g_p(\mathbf{r}) + \left(1-g_p(\mathbf{r})\right)\ln\left(1-g_p(\mathbf{r})\right) - \frac{g_p(\mathbf{r})}{2}\left(\ln\left(v_p\left\langle n_p(\mathbf{r})\right\rangle\right)-\beta\varepsilon_p\right)\right] \\
& + \sum_{p\in \mathrm{FG,NQT}}\int d\mathbf{r}\frac{1}{v_p}\left[1-\left(1-\frac{g_p(\mathbf{r})}{2}\right)v_p\left\langle n_p(\mathbf{r})\right\rangle\right]\ln\left[1-\left(1-\frac{g_p(\mathbf{r})}{2}\right)v_p\left\langle n_p(\mathbf{r})\right\rangle\right]
\end{aligned} \tag{S1}
$$

where $\beta = 1/k_B\mathrm{T}$ and $\mathbf{r}$ is the position vector.

The first two terms in Eq. S1 represent translational (mixing) entropies of solvent molecules, cations, anions, protons and hydroxyl ions, where $\rho_i(\mathbf{r})$ the number of species *i* at $\mathbf{r}$ (i = A, C, w, $H^+$, $OH^-$ for anions, cations, solvent molecules, protons and hydroxyl ions) and $\mu_i^0$ is the standard chemical potential of species *i*. The third term is the conformational entropy of the disordered protein chains, where $\mathbf{s}$ is a parameterization of the membrane area with tethered chains, d$\mathbf{s}$' is the area element and the integral runs over the area where the FG-Nups are anchored, $\sigma_k(\mathbf{r})$ is the grafting density of chains of type *k* at $\mathbf{r}$, $P_k(\mathbf{r}, \alpha)$ is the probability of having a chain of type *k* anchored at $\mathbf{r}$ in conformation $\alpha$ and $N_k$ is the number of different FG-Nups in the system. In this term, d$\mathbf{s}$ is the corresponding area element. The fourth term is the effective attraction between amino-acid beads, which represents the difference between the segment-segment and segment-solvent vdW attraction energies. In this term, $\langle n_l(\mathbf{r})\rangle$ is the number density of the amino acid of type *l* ($N_l$ is the number of different amino acids) at $\mathbf{r}$:

$$\left\langle n_l(\mathbf{r})\right\rangle = \int d\mathbf{s}' \sum_k \sigma_k\left(\mathbf{r}'(\mathbf{s}')\right)\sum_\alpha P_k\left(\mathbf{r}'(\mathbf{s}'),\alpha\right) n_{kl}\left(\mathbf{r}'(\mathbf{s}'),\alpha,\mathbf{r}\right) \quad \text{(S2)}$$

In Eq. S2, $n_{kl}(\mathbf{r}'(\mathbf{s}'), \alpha, \mathbf{r})d\mathbf{r}$ is the number of amino acids segments of type $l$ that a protein of type $k$ tethered at $\mathbf{r}'(\mathbf{s}')$ has in the volume element between $\mathbf{r}$ and $\mathbf{r} + d\mathbf{r}$ when it is in conformation $\alpha$. The expression $g(|\mathbf{r} - \mathbf{r}'|)$ is a distance-dependent vdW attractive interaction of the form:

$$g\left(\left|\mathbf{r}-\mathbf{r}'\right|\right) = \begin{cases} -\left(\dfrac{a}{\left|\mathbf{r}-\mathbf{r}'\right|}\right)^6 & a < \left|\mathbf{r}-\mathbf{r}'\right| < 1.5\delta \\ 0 & \text{otherwise} \end{cases} \quad \text{(S3)}$$

where $a$ is the segment length and $1.5\delta$ is a cut-off parameter. The parameter $\varepsilon_{ll'}$ (in $k_BT$ units) determines the strength of the interaction between amino acids of type $l$ and $l'$. The fifth term in Eq. S1 is the electrostatic contribution to the free energy, where $\psi(\mathbf{r})$ is the local electrostatic potential, $\varepsilon$ is the dielectric constant and $<\rho_Q(\mathbf{r})>$ is the average charge density at $\mathbf{r}$, given by

$$\left\langle \rho_Q(\mathbf{r})\right\rangle = \sum_{i\in \mathrm{A,C,H^+,OH^-}} \rho_i(\mathbf{r})q_i + \sum_l \left\langle n_l(\mathbf{r})\right\rangle f_l(\mathbf{r})q_l \quad \text{(S4)}$$

where the sum in the first term on the RHS runs over all charged mobile species, $q_l$ is the nominal charge of the side chain of an amino acid of type $l$ (i.e. -1 for acidic amino acids and +1 for basic amino acids), $f_l$ ($\mathbf{r}$) is the fraction of charged amino acids of type $l$ that are charged at $\mathbf{r}$ (the summation runs only over chargeable amino acids).

The last two terms in Eq. S1 describe the pairing interactions between stickers (amino acids that have the potential to form a hydrogel), which include both FG motifs and cohesive spacers NQT. In these terms, $\varepsilon_p$ describes the effective interaction strength (including the solvent effect) between the stickers, and $v_p$ is the effective volume of the pair (the volume within which the stickers will pair up). The pairing fraction $g_p$ varies in the range from 0 to 1 describes how saturated the gelation is. Note that the pairing interaction depends on both the pair strength and volume, and thus is more than a reaction between different internal states (dangling and paired) of the stickers even in a mean-field language. This mean-field treatment of gelation is qualitatively different from our previous model where a vdW form (Eq. S2) was used regardless of the nature of the cohesive interaction(Tagliazucchi et al., 2013).

The system's free energy (Eq. S1) is minimized with the constraint of the incompressibility assumption of the system, which accounts for the excluded volume repulsion:

$$\sum_{i\in \mathrm{A,C,w,H^+,OH^-}} \rho_i(\mathbf{r})v_i + \sum_{l=1,N_l} v_l \left\langle n_l(\mathbf{r}) \right\rangle = 1 \tag{S5}$$

where $v_i$ is the volume of the species $i$ (anions, cations, water molecules, protons, hydroxyl ions) and $v_l$ is the volume of polymer monomers of type $l$. This packing constraint is enforced by introducing the Langrage multiplier, which is a position-dependent osmotic pressure.

The details of the free energy functional minimization have been documented in our previous works(Tagliazucchi et al., 2013). For the new pairing term, minimization leads to an analytical form for the pairing fraction:

$$g_p(\mathbf{r}) = 1 - \frac{1}{2\exp(-\beta\varepsilon_p)-1}\left[\left(2\exp(-\beta\varepsilon_p)-1+\frac{1}{v_p\left\langle n_p(\mathbf{r})\right\rangle}\right)^2 - 4\exp(-\beta\varepsilon_p)\left(\exp(-\beta\varepsilon_p)-\frac{1}{2}\right)\right]^{\frac{1}{2}} + \frac{1}{v_p\left\langle n_p(\mathbf{r})\right\rangle\left(2\exp(-\beta\varepsilon_p)-1\right)} \tag{S6}$$

which is a scalar field in space as it depends on the local density $n_p$ of the stickers.

### NPC model

We grouped the amino acids into 10 categories according to their properties. Different groups of amino acids used in our NPC model are summarized in Table S1. The listed pKa values are for dilute conditions, and effect of local pKa shift has been explicitly considered in our model. We consider the volume of all amino acids to be equal to the average value of 0.095 nm$^3$.

**Table S1: Classification of amino acids into different groups.**

| Group (*l*) | Amino acid code* | Acid-base properties | Solubility property |
|---|---|---|---|
| 1 | Fx | Neutral | Hydrophobic |
| 2 | xFOx | Neutral | Hydrophobic |
| 3 | FxOx | Neutral | Hydrophobic |
| 4 | N, Q, T | Neutral | Hydrophilic, cohesive |
| 5 | A, I, L, W, Y | Neutral | Hydrophobic |
| 6 | G, M, P, S, V | Neutral | Hydrophilic |
| 7 | K, R | Base, pKa = 11 | Hydrophilic |
| 8 | D, E | Acid, pKa = 5 | Hydrophilic |
| 9 | C | Acid, pKa = 8.3 | Hydrophilic |
| 10 | H | Base, pKa = 6.08 | Hydrophilic |

* x means hydrophilic, O means hydrophobic. For FG motifs such as xFOx, the group ID is assigned to F. If there are two F in the same FG motifs, one of them will be classified as F stickers ($l$=1,2,3), the other assigned to $l$=5.

Besides steric and electrostatic interactions, we considered two kinds of cohesive interactions, namely the pairing interaction and the vdW interaction. All the FG motifs are assigned the same pairing energy and can associate with other hydrophobic amino acids through vdW interaction. Between the FG motifs, vdW interaction is excluded (for amino acids of low concentration, the vdW interaction is negligible in any case). While all the FG motifs have the same pairing energy regardless their types, type-2 and type-3 FG motifs have an extra hydrophobic amino acid that results in slightly more cohesiveness (from vdW interactions). Due to the abundance of the cohesive spacers (NQT) and their propensity to form $\beta$ -sheet, we assigned both vdW and pairing interactions to them. Table S2 summarizes the interaction coefficients between all amino acids pairs for a reasonable level of cohesion based on which most of our analyses in the main text are done. The first value corresponds to the pairing energy, the second to the vdW energy. We chose the effective pairing volume for FG pairs to be 20 times the average volume of an amino acid due the relatively long-range interaction between FG pairs, and 6 times for the pairing volume for spacer pairs. To illustrate the cooperation between the FG motifs and the cohesive spacers, we have repeated our calculations for different combinations of the pairing and vdW energies and shown the results in Fig. 1 in the main text. Besides interactions between amino acids, we have assigned a weak interaction (0.15kT) between hydrophobic/cohesive amino acids and the NPC scaffold surface (of the inner ring), in accord with the recent experimental observations of scaffold cohesiveness.

**Table S2: Values of the interaction coefficients $\varepsilon_p$*, $\varepsilon_{ll'}$ in the units of kT**

| | ***l*=1-3** | ***l*=4** | ***l*=5** | ***l*=6-10** |
|---|---|---|---|---|
| ***l*=1-3** | 2.5, 0 | 0, 0 | 0, 2 | 0, 0 |
| ***l*=4** | 0, 0 | 1, 1 | 0, 0 | 0, 0 |
| ***l*=5** | 0, 2 | 0, 0 | 0, 2 | 0, 0 |
| ***l*=6-10** | 0, 0 | 0, 0 | 0, 0 | 0, 0 |

* $\varepsilon_p$ only exists for diagonal elements as pairing happens between the same category of stickers

We model the FG-Nups as freely jointed linear chains made of $N$ hard sphere monomers at fixed bond length. We chose the sequences and grafting positions of the FG-Nups (see Figure 1 in the main text and table S3) based on the data from the recent work of Rout and coworkers(Kim et al., 2018). In this work, the stoichiometry of NPC (the copy numbers of different FG-Nups) was found to be nearly doubled compared to the previous experiments(Alber et al., 2007). The parts that connect between the folded and unfolded subdomains of the FG-Nups lack the atomic resolution of the scaffold and therefore lead to uncertainties in the exact anchoring positions both in terms of sequence and in space. Nevertheless, uncertainties of a few nanometers in space and tens of amino acids for the anchors are not expected to affect the main conclusions of our work. The properties of the IDRs, their stoichiometry and anchoring positions are summarized in Tables S3 and S4. The lengths of IDRs are in number of amino acids. Compared to our previous model(Tagliazucchi et al., 2013), we excluded the short FG-poor Nup53 and Nup59 IDRs and added the long FG-rich Nup2 IDR.

**Table S3: Properties of the IDRs considered in the model.**

| FG-Nup | length | domain | *l*=1 | *l*=2 | *l*=3 | *l*=4 | *l*=5 | *l*=6 | *l*=7 | *l*=8 | *l*=9 | *l*=10 |
|---|---|---|---|---|---|---|---|---|---|---|---|---|
| Nup159 | 695 | 388-1082 | 24 | 15 | 3 | 114 | 94 | 269 | 63 | 103 | 0 | 10 |
| Nup100 | 801 | 1-800 | 19 | 33 | 1 | 273 | 108 | 298 | 39 | 26 | 0 | 4 |
| Nup116 | 966 | 1-966 | 21 | 45 | 1 | 320 | 150 | 327 | 55 | 44 | 1 | 2 |
| Nup42 | 382 | 1-382 | 19 | 12 | 2 | 118 | 57 | 159 | 13 | 2 | 0 | 0 |
| Nup49 | 270 | 1-270 | 3 | 13 | 1 | 89 | 48 | 106 | 8 | 1 | 1 | 0 |
| Nsp1 | 601 | 1-601 | 8 | 7 | 21 | 133 | 103 | 196 | 71 | 62 | 0 | 0 |
| Nup57 | 287 | 1-287 | 3 | 10 | 3 | 109 | 42 | 113 | 7 | 0 | 0 | 0 |
| Nup145N | 426 | 1-426 | 5 | 16 | 1 | 113 | 78 | 159 | 31 | 22 | 0 | 1 |
| Nup1* | 876 | 1076-201 | 11 | 4 | 20 | 186 | 151 | 290 | 115 | 90 | 2 | 7 |
| Nup2 | 720 | 1-720 | 11 | 6 | 14 | 129 | 130 | 220 | 104 | 104 | 1 | 1 |
| Nup60* | 189 | 539-351 | 1 | 4 | 0 | 32 | 39 | 61 | 26 | 22 | 0 | 4 |

* takes a reversed direction from anchor to free end compared to the other FG-Nups

**Table S4: Stoichiometry and anchoring positions of the FG-Nups.**

| FG-Nup | Stoichiometry | Anchoring r (nm) | Anchoring z (nm) |
|---|---|---|---|
| Nup159 | 16 | 36.8 | 13.5 |
| Nup100 | 16 | 33.1, 35.2 | 7.4, 13.4 |
| Nup116 | 16 | 29.0, 36.2 | 18.1, 21.8 |
| Nup42* | 8 | 31 | 13 |
| Nup49 | 32 | 22.9, 21.1, 22.9, 21.1 | 3.0, 3.4, -3.0, -3.4 |
| Nsp1 | 48 | 28.0, 31.0, 23.2, 23.0, 23.2, 23.0 | 16.0, 18.4, 4.9, 4.5, -4.9, -4.5 |
| Nup57 | 32 | 23.9, 22.0,23.9, 22.0 | 1.6,2.1,-1.6,-2.1 |
| Nup145N | 16 | 35.2, 30.7 | -12.4,-7.2 |
| Nup1* | 16 | 35 | -9.1 |
| Nup2* | 16 | 31 | -22 |
| Nup60 | 16 | 37.2, 36.1 | -16.5,-19.3 |

* FG-Nups that lack well characterized stoichiometry or anchoring positions.

To account for the conformational entropy of the IDRs, we generated a large set of chain conformations (~$10^8$ in total for all the IDRs). The time required for such chain generation is computationally prohibitive using a simple generation routine due to the amino acid-scaffold and intrachain amino acid-amino acid excluded volume requirements; therefore, we have implemented an enriched sampling method(Grassberger, 1997). More details about chain generation, symmetry considerations, discretization and numerical solution can be found in our previous paper(Tagliazucchi et al., 2013).

### FG-pairing energy

The FG-pairing energy largely determines the propensity of gelation for the FG-Nups. To estimate this important energetic parameter, we carried out molecular dynamics (MD) simulations with explicit water molecules to capture the hydrophobic effect. The interaction energy profile between two phenylalanine side chains, shown in Fig. S1, was obtained from a series of 20 all-atom MD simulations. Each simulation has two phenylalanine monomers placed at the center of a 10 nm x 10 nm x 10nm cubic box filled with ~33000 explicit water molecules. Across the 20 simulations the distance between two backbone C-alpha atoms was systematically varied from 0.1 nm to 2.0 nm, with an interval of 0.1 nm. These C-alpha distances were held fixed during the simulations, while all the other atoms of the phenylalanine molecules as well as the water molecules were allowed to move freely. The OPLS force field(Jorgensen et al., 1996) was used for phenylalanine, and the TIP4P potential(Jorgensen and Madura, 1985) was used for water. The temperature was held at 300K via the Nosé-Hoover thermostat(Nosé, 1984) during the simulations, and the volume of the simulation box was kept constant. After the completion of all 20 simulations of 10 ns each, the interaction energy between the two phenylalanine side chains was extracted to create the interaction energy profile. All simulations were run on the GROMACS simulation package(Pronk et al., 2013), version 4.6.1.

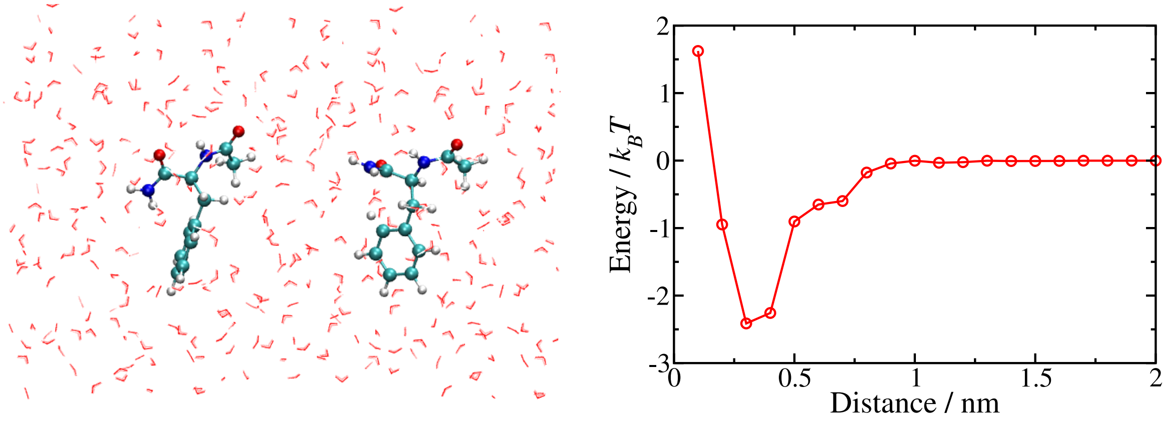


**Figure S1. FG-pairing energy calculated from all-atom MD simulation.** Left: two phenylalanine molecules in explicit water (showing only part of the simulation box). Right: the interaction energy profile of the two side chains of the amino acids.

### Supplemental analyses

In Figs. 2D, F of the main text we showed the spatial distribution of net charge from the IDRs and the DEKR spacers (total charge) distribution. The two distributions differ widely in both magnitude and shape, with the latter being denser and more heterogeneous. This is because the positive and negative amino acids have similar spatial distributions that largely cancel out, as shown in Figs. S2A, B. It is remarkable that despite the heterogeneous distributions of both the positive and negative amino acids, the resultant net charge distribution appears relatively homogeneous (Fig. 2D). In Fig. 2C of the main text we showed that the spatial distribution of the cohesive spacers (NQT) is segregated from that of the charged amino acids. In contrast, the spatial distribution of non-cohesive neutral spacers is less anti-correlated with the charged spacers, as shown in Fig. S2C. The spatial distribution of the non-Phenylalanine hydrophobic amino acids is shown in Fig. S2D.

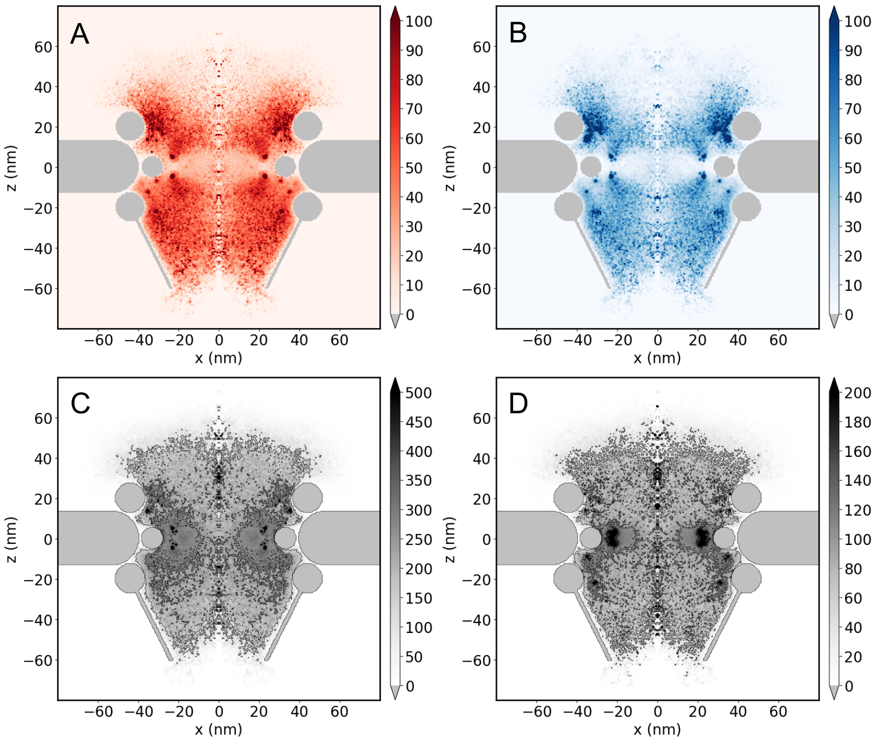


**Figure S2. Spatial distributions of some amino-acid groups in mM concentration.** (**A**) Positive amino acids. (**B**) Negative amino acids. (**C**) Non-cohesive neutral spacers. (**D**) Non-Phenylalanine hydrophobic amino acids

The morphology of the thermoreversible FG network, despite its low pairing fraction, is more structured than that of a non-cohesive brush. To see this more clearly, we plot the morphology, FG distribution and pairing fraction map of a non-cohesive NPC in Fig. S3. In this system, both the FG pairing interaction and the attraction between NQT spacers are turned off. As shown in Fig. S3A, the brush morphology lacks the condensed central barrier ring and the vestibular structures at the exits of the pore, compared to colored panel in Fig. 1C. The FG motifs are widely distributed throughout the NPC (Fig. S3B) and the pairing fraction is just a few percent throughout the pore as shown in Fig. S3C. It is non-zero since there is a chance of two non-sticky FG motifs getting into contact by thermal fluctuations. Compared to the cohesive case, the non-cohesive FG-Nups are more spread out as shown in Fig. S4. The non-cohesive Nup116, Nup100 can no longer close the cytoplasmic side of the NPC.

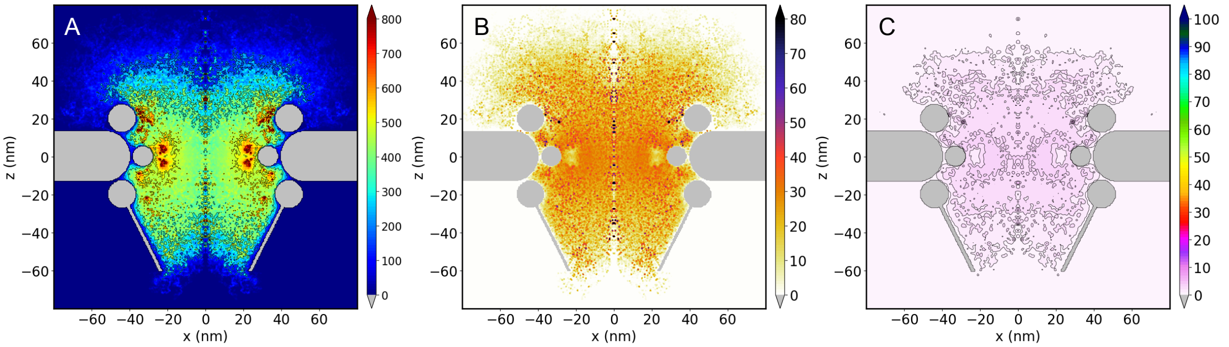


**Figure S3. Non-cohesive brush system.** (**A**) Overall IDR concentration in mM. (**B**) FG spatial distribution in mM. (**C**) FG-pairing fraction in percentage.

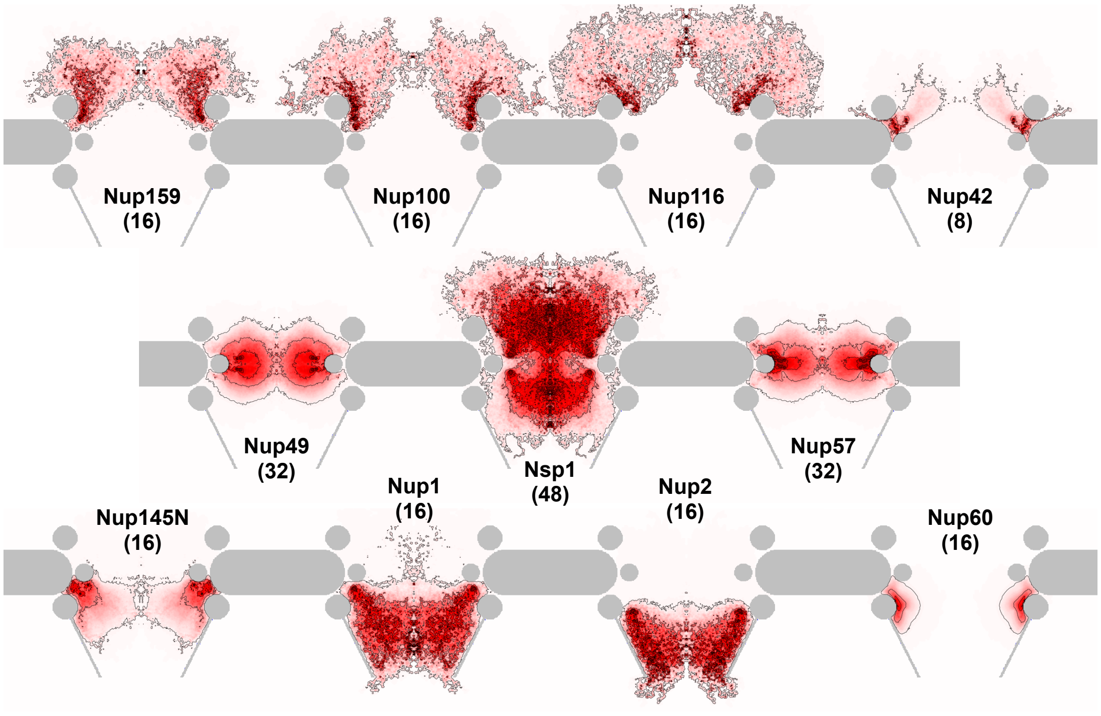


**Figure S4. Spatial distributions of non-cohesive FG-Nups.**

Lastly, it is instructive to show how the FG pairing and the attraction between cohesive spacers tend to shape the central transporter of NPC differently. To this end we modeled two extreme cases. In the first one we assigned strong FG pairing interaction (4.5kT) and turned off the attraction between cohesive spacers. In contrast, we used highly cohesive spacers (1.4kT attraction) and non-sticky FG motifs in the second system. The overall morphologies, distributions of FG motifs and their pairing fraction maps are shown in Fig. S5 for both cases. In the FG-dominant system, we found a considerably homogenous gating morphology (Fig. S5A) with relatively uniform spatial distribution of the FG motifs (Fig. S5B) and high pairing fraction in the pore center (above 50%, Fig. S5C). This prediction suggests that FG pairing alone tends to form a homogeneous hydrogel. Nevertheless, such gelation requires a pairing strength that is much higher than the thermal energy, which would make the FG network less thermoreversible than suggested by experiments(Hough et al., 2015). In the spacer-dominant system, the condensation of the cohesive spacers leads a highly heterogeneous gating structure (Fig. S5D), whose pattern is to some degree similar to what we have predicted under reasonably cohesive conditions (main text and Table S2). This result reinforces our conclusion that the cohesive spacers are crucial in guiding the self-assembly of the central transporters. The heterogeneous gating structure is accompanied by an inhomogeneous spatial distribution of FG motifs (Fig. S5E). However, without affinity, the FG pairing fraction is very low, as shown in Fig. S5F. The phase separation between high and low density regions is in stark contrast to the gelation in the first system.

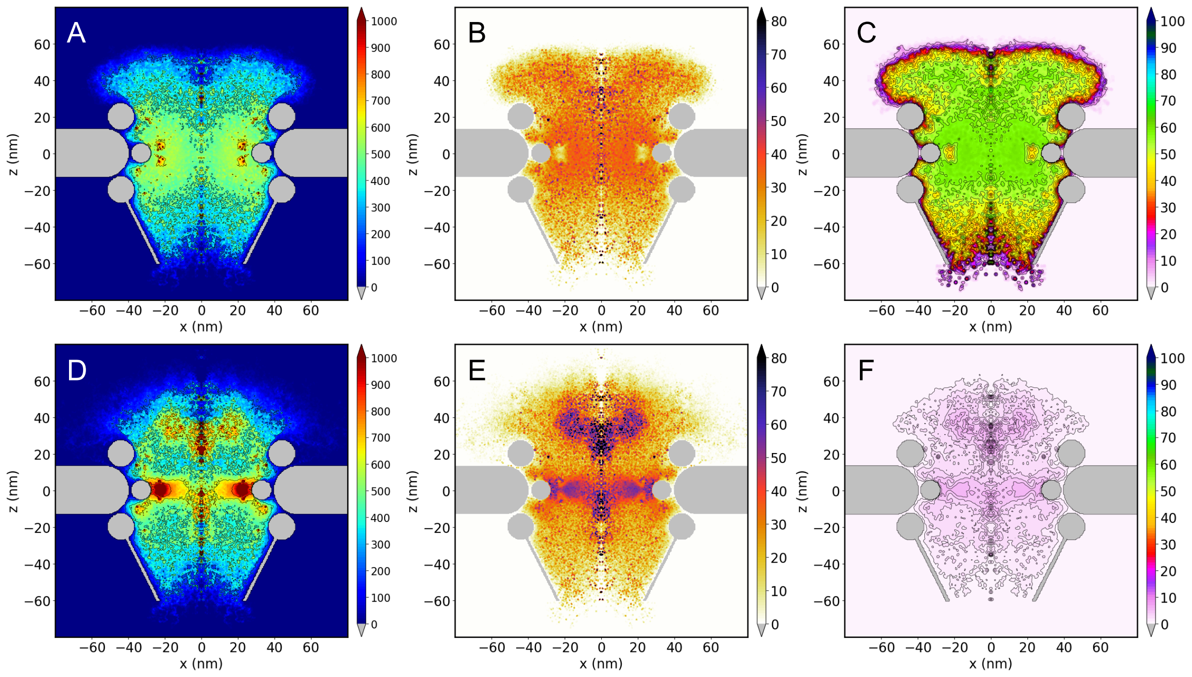

**Figure S5. Comparison between two extreme models of NPC.** (**A-C**) Overall morphology of the FG-Nups, spatial distribution of FG motifs and their pairing fraction map for a hydrogel system. (**D-F**) Same analyses for the system with phase separation. (Panels A, B, D, E are in mM and C, F in percentage.)
